# Copy number of the eukaryotic rRNA gene can be counted comprehensively

**DOI:** 10.1101/2023.08.18.553942

**Authors:** Akinori Yabuki, Tatsuhiko Hoshino, Tamiko Nakamura, Keiko Mizuno

## Abstract

Gene sequence has been widely used in molecular ecology. For instance, the ribosomal RNA (rRNA) gene has been widely used as a biological marker to understand microbial communities. The variety of the detected rRNA gene sequences reflects the diversity of the microorganisms existing in the analyzed sample. Their biomass also can be estimated by applying quantitative sequencing with the information of rRNA gene copy numbers in genomes, however, the information on rRNA gene copy number is still limited. Especially, the copy number in microbial eukaryotes is much less understood than that of prokaryotes, possibly because of the large and complex structure of eukaryotic genomes. In this communication, we report an alternative approach that is more appropriate from the existing method of quantitative sequencing and demonstrate that the copy number of eukaryotic rRNA can be measured efficiently and comprehensively. By applying this approach widely, information on the eukaryotic rRNA copy number can be determined, and their community structures can be depicted and compared more efficiently.

## Main Text

The ribosomal RNA (rRNA) gene encodes a fundamental component of the ribosome, and its sequence information has been utilized in a very wide range of biological studies. For instance, the 18S rRNA gene (i.e., the eukaryotic small subunit rRNA gene) is most frequently used as a biological marker. The existence and distribution of microbial eukaryotes have been analyzed based on 18S rRNA gene sequences in environmental analyses (e.g., de Vargas *et al*. [1]). The 18S rRNA genes of many microbial eukaryotes have been sequenced and deposited in GenBank. Although the copy numbers of prokaryotic rRNA genes are well understood and summarized in *rrn*DB [2], it is much less known about their copy numbers in eukaryotic genomes. The copy number of the rRNA gene is important for addressing systematic bias when measuring community composition in molecular surveys based on rRNA gene abundance, and it helps provide a robust evaluation of the diversity and distribution of eukaryotes in the natural environment. Notably, the copy number has been counted in several protists using various approaches [3]; some were estimated by quantitative PCRs [4], and others were directly counted using the genomic information [5]. These approaches are sufficiently operational but time- and effort-consuming. Therefore, a more efficient approach is required. Quantitative sequencing (qSeq) allows the simultaneous sequencing of amplicons and comprehensive gene quantification [6,7], and the unique sequence tags introduced by the single primer extension at the first step at the end of a target DNA molecule are counted after sequencing to estimate the abundance of the target DNA. In the present study, we demonstrated that qSeq can be used to efficiently and comprehensively quantify eukaryotic rRNA genes.

We analyzed five diplonemid species whose community structure and exchanges in natural environments have recently been focused on because this information may be useful for monitoring biodiversity in certain environments [8]. The genomic DNA of each deplonemid species was extracted after counting the cells (Table 1). Simultaneous quantification and sequencing of diplonemid DNA were performed as previously described [9], with a few modifications. Briefly, after single primer extension using S616F_Cerco primer [10] at 68 °C for 10 min, 8 μl of ExoSAP-IT Express was added to 20 μl of single primer (SPE) reaction to digest excess primers, followed by incubation at 37 °C for 4 min. The SPE product was used for the first round of PCR with the S948_Dip primer [11] and an adapter sequence for the second round of PCR. The PCR product was purified using agarose gel electrophoresis, followed by an index PCR for sequencing. After purification using AMpure XP beads, the indexed PCR products were sequenced using a MiSeq platform with the MiSeq Reagent Kit v3 for 600 cycles (Illumina). Quality trimming and merging of sequence reads were performed using Mothur ver.1.44 [12]. Unique DNA tags were counted as described previously [13]. The experimental process is summarized schematically in Figure 1A.

**Table 1.**

| Sample | Cell number<br>for DNA extraction | DNA density<br>(ng/μl) | Abundance<br>(copies/μl of DNA) | Copy number (copies/cell) |  |  |
| --- | --- | --- | --- | --- | --- | --- |
|  |  |  |  | Each<br>replicate | Average | Median |
| *Diplonema papillatum* -1 | 1.00E+06 | 1.22 | 1.7E+05 | 17.2 | 17.8 | 17.2 |
| *Diplonema papillatum* -2 | 1.00E+06 | 0.94 | 1.6E+05 | 15.6 |  |  |
| *Diplonema papillatum* -3 | 1.00E+06 | 1.06 | 2.1E+05 | 20.5 |  |  |
| *Rhynchopus* sp. YPF1506 -1 | 3.00E+05 | 0.32 | 3.2E+04 | 10.8 | 7.0 | 5.1 |
| *Rhynchopus* sp. YPF1506 -2 | 3.00E+05 | 0.21 | 1.5E+04 | 5.1 |  |  |
| *Rhynchopus* sp. YPF1506 -3 | 3.00E+05 | 0.23 | 1.5E+04 | 5.0 |  |  |
| *Lacrimia lanifica* -1 | 3.00E+05 | 0.21 | 6.2E+04 | 20.5 | 23.4 | 20.5 |
| *Lacrimia lanifica* -2 | 3.00E+05 | 0.21 | 6.0E+04 | 19.9 |  |  |
| *Lacrimia lanifica* -3 | 3.00E+05 | 0.26 | 9.0E+04 | 29.9 |  |  |
| *Sulcionema specki* -1 | 1.00E+05 | 0.98 | 7.7E+04 | 76.6 | 73.2 | 76.6 |
| *Sulcionema specki* -2 | 1.00E+05 | 1.21 | 8.2E+04 | 82.3 |  |  |
| *Sulcionema specki* -3 | 1.00E+05 | 0.73 | 6.1E+04 | 60.6 |  |  |
| *Hemistasia phaeocysticola* -1 | 3.00E+05 | 0.08 | 6.9E+03 | 2.3 | 1.8 | 1.9 |
| *Hemistasia phaeocysticola* -2 | 3.00E+05 | 0.077 | 5.7E+03 | 1.9 |  |  |
| *Hemistasia phaeocysticola* -3 | 3.00E+05 | 0.072 | 3.4E+03 | 1.1 |  |  |

**Fig. 1 A.**
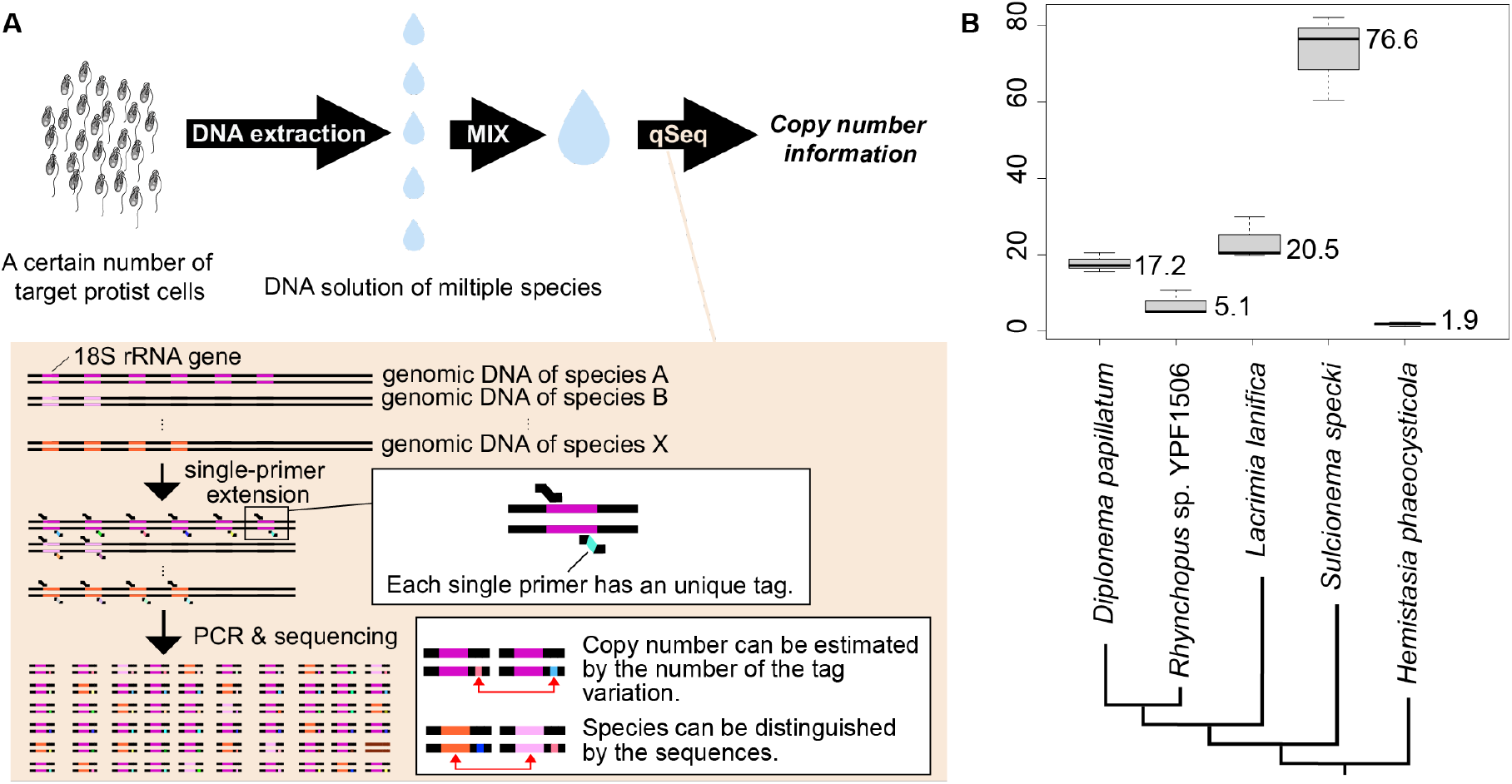
Schematic image of the experimental process to comprehensively count the copy number of the eukaryotic rRNA gene. The more detailed process of qSeq was summarized in Hoshino and Inagaki [6]. B. The result of the rRNA gene copy number estimation of five diplonemids. The value next to the boxes is the median of each species. The phylogenetic relationships among the five species were referred from Tashyreva *et al*. [15]

The obtained copy number of the 18S rRNA gene of each species from the three replicates was consistent (Table 1; Fig. 1B); however, it differed among the species: the largest was approximately 77 (*Sulcionema specki*) and the smallest was approximately 2 (*Hemistasia phaeocysticola)*. As *Diplonema papillatum*, whose rRNA gene copy number has already been counted from genomic sequence data [14], has approximately 21 copies, our estimated score of *D. papillatum* (i.e., approximately 17) is comparable to it, and the estimation of the other four species can be expected to be reasonable. Minor inconsistencies in the estimated copy numbers of *D. papillatum* may be due to technical errors in cell counting and/or DNA extraction. With qSeq, the efficiency of the first round of PCR does not affect the quantitative results but can cause underestimation if the efficiency of the first SPE extension is not 100%. However, if exonuclease treatment after SPE is insufficient, the primer may remain, leading to overestimation [9]. Nevertheless, the impact of possible artifacts in the experimental process was minor, and our estimation is reasonable. Although rRNA gene copy numbers have never been compared among diplonemids, this study revealed some variation. All diplonemids have similar lifestyles (i.e., planktonic flagellates feeding on microorganisms, including zooplankton and algae) and similar cell sizes; however, their rRNA gene expression may be regulated in different ways. Their copy numbers increased and/or decreased multiple times during evolution because they did not simply increase or decrease along with their phylogeny (Fig. 2B). This variation is important when discussing the distribution and community structure of diplonemids. Species with higher copy numbers can be detected more easily using environmental DNA analysis, and the number of detected sequences does not simply correspond to their biomass. In the present study, we showed that qSeq can accurately quantify the rRNA gene copy numbers of five diplonemid species, indicating that it can be applied to other eukaryotic microorganisms for which the rRNA copy number is unknown. In particular, the ability of qSeq to simultaneously quantify the copy number of genes in many species is advantageous for analyses in natural environments. The scale of the analysis, that is, the number of analyzed species and the cell number for DNA extraction, was also flexible in the present approach; the DNA of each diplonemid species was extracted from 1, 3, or 10E+05 cells in the present study. Notably, quantification of rRNA gene copy numbers in individual eukaryotic microorganisms in the environment using qSeq can lead to a further understanding of their ecology and diversity.

## Acknowledgements

This study was supported in part by the Japan Society for the Promotion of Science (20K06792 to A.Y. and 21K19876 to T.H.).

## Data Availability Statement

The raw sequencing data for copy number estimation are available under GenBank BioProject accession number SSUB026492.

## Author Contributions

Conceptualization: A.Y. & T.H.; Cell measurements and DNA preparation: A.Y., K.M., T.N.; Sequencing and data analyses: T.H.; Original draft and Writing: A.Y., T.H. All authors have read and confirmed the final version of the manuscript.

